# Fibrin Prestress Due to Platelet Aggregation and Contraction Increases Clot Stiffness

**DOI:** 10.1101/2021.04.19.440330

**Authors:** Suyog J. Pathare, Wilson Eng, Sang-Joon J. Lee, Anand K. Ramasubramanian

**Author notes:** **Corresponding authors** Sang-Joon J. Lee, Anand K. Ramasubramanian, **Emails:**.

## Abstract

Efficient haemorrhagic control is attained through the formation of strong and stable blood clots at the site of injury. Although it is known that platelet-driven contraction can dramatically influence clot stiffness, the underlying mechanisms by which platelets assist fibrin networks in resisting external loads are not understood. In this study, we delineate the contribution of platelet-fibrin interactions to clot tensile mechanics using a combination of new mechanical measurements, image analysis, and structural mechanics simulation. Based on uniaxial tensile test data using custom-made microtensometer, and fluorescence microscopy of platelet aggregation and platelet-fibrin interactions, we show that integrin-mediated platelet aggregation and actomyosin-driven platelet contraction synergistically increase the elastic modulus of the clots. We demonstrate that the mechanical and geometric response of an active contraction model of platelet aggregates compacting vicinal fibrin is consistent with the experimental data. The model suggests that platelet contraction induces prestress in fibrin fibres, and increases the effective stiffness in both crosslinked and non-crosslinked clots. Our results provide evidence for fibrin compaction at discrete nodes as a major determinant of mechanical response to applied loads.

## 1. Introduction

Efficient haemostasis culminates in the formation of strong and stable plugs to dam blood flow in broken vessels or to stop leaks through injured vessel walls. The strength and stability of clots are important predictors of bleeding diathesis and thrombotic complications, and these properties are being adopted for new indications including trauma and anticoagulant monitoring (1,2). Clot components including platelets, fibrinogen, and red blood cells contribute to the mechanical properties of the clot (3–6). The dynamics of interactions between these components on clot mechanics is an active area of investigation because of fundamental importance and translational benefits.

The fibrin network is the key structural element of a clot. Platelets not only initiate and propagate fibrin network formation but also modify the structure of the fibrin network extensively during the later stages of clotting through retraction. The haemostatic potential of the platelets including their ability to activate, to aggregate, and--most importantly--to retract determines the clot stiffness (7–9). It was recently shown that clot contraction and enhanced clot stiffness are the results of exquisitely organized, platelet-driven, actomyosin- and integrin-mediated remodelling of fibrin networks (10). However, the mechanism by which platelet contraction enhances resistance of a clot network against applied forces is unknown, and this question is of fundamental importance to the understanding of haemostasis.

To answer this question, first we evaluate the stress-strain behaviour of blood clots to uniaxial tensile strain using a custom-made microtensometer. To our knowledge, this is the first study to report the elastic properties of human blood clots subjected to tensile strain. Uniaxial testing is a direct method to measure the tensile mechanical properties of several (vascular) biomaterials, including fibrin gels (11), collagen gels (12), cell-seeded scaffolds (13), and aortic tissue (14). Together with appropriate mathematical models, tensile tests have provided profound insights into the mechanobiology of cell-matrix interactions (15–18). Second, using the experimental tensile and microstructural data in conjunction with a computational model for fibrin network response to platelet compaction strain and externally applied strain, we elucidate the relationship between platelet-fibrin interactions and the mechanical response of clots to applied extension. We discover that platelet aggregates compacted with dense fibrin bundles act as prestressed nodes that dramatically increase the stiffness of the entire clot network.

## 2. Materials and Methods

### 2.1. Measurement of clot response to uniaxial strain

Blood was obtained from healthy adult donor volunteers in 3.2% sodium citrate, and clotting was initiated within 15 min of blood draw after recalcification with 20 mM CaCl_2_ (Sigma, St. Louis, MO, USA). The clotting process was initiated by the addition of thrombin to 500 µL of recalcified blood, and the clot was incubated under humidified conditions for 1.5 h for full contraction. The clots were gently separated from the serum and loaded onto the clamps of a custom made microtensometer. The microtensometer consists of a motorized nanopositioner (MP-285, Sutter Instrument, Novato, CA, USA), a full-bridge thin cantilever load cell (113 g, LCL Series, OMEGA Engineering, Norwalk, CT, USA), and a data acquisition (DAQ) signal conditioner DI-1000U (Loadstar Sensors, Fremont, CA, USA). The load cell was zeroed without the bottom support to neglect the weight of the clot in the initial pull. The force recording from the data acquisition software (LoadVUE, Loadstar Sensors, Fremont, CA, USA), and actuation of the nanopositioner were started simultaneously, and force measurements were recorded at 30 Hz. Two digital cameras were positioned at orthogonal angles to capture top-view and side-view images to extract the cross-sectional dimensions. Cross-section images were used to convert force-displacement curves to the respective true stress vs. stretch ratio curves.

In designated cases, the following inhibitors were added to blood, and incubated for 30 min at room temperature with gentle rocking (Vari-Mix Platform Rocker, Thermo Fisher Scientific, Waltham, MA, USA) at approximately 2 Hz before recalcification: 300 μM blebbistatin; 10 µM eptifibatide; or 10 µM T101. To investigate the effect of platelets in clotting, red blood cells (RBCs) were separated by centrifugation for 15 min without brakes at 250 relative centrifugal force (RCF), 7 rad/s^2^ acceleration, and after separating the RBCs from platelet rich plasma (PRP), platelet poor plasma (PPP) was obtained by further centrifugation of PRP for 20 min without brakes at 1500 RCF, 7 rad/s^2^ acceleration. The RBC was added back to PPP to restore the haematocrit to original levels in the whole blood.

### 2.2. Microstructural characterization of clots

To visualize clot microstructure, platelets and fibrinogen were labelled with AF488 and AF596 fluorophores, and 100 μL of blood was used to form clots as described above. Platelets were incubated with AF488-conjugated CD42b antibody for 20 min prior to the start of clotting, and blood was doped with 5% AF596 conjugated fibrinogen assuming a final concentration of 4 mg/mL. After 1.5 h, the clots were imaged at 63x magnification using an oil immersion lens (DMi8, Leica Microsystems, Germany). Z-stack images were collected from at least five different locations for each clot. The fluorescence images were processed and analyse offline for quantification by 2D projection of the stack to estimate aggregate size and Pearson Correlation Coefficient (PCC) using Image-Pro 10 (Media Cybernetics, Rockville, MD, USA) and ImageJ (NIH).

### 2.3. Discrete prestressed fibre model simulations

Simulations were performed using COMSOL Multiphysics (Stockholm, Sweden), using quasi-static 2-D truss elements with geometric nonlinearity enabled. Compaction strain *α* was mimicked using a thermal expansion coefficient. Convergence of the discrete fibre model was checked by varying the number of fibres and evaluating the average strain energy density. The range of solution values among five-fibre, six-fibre, and seven-fibre solutions was less than 0.6% of the mean, and five fibres were used for all reported strain energy density simulations.

### 2.4. Statistical analysis

The data was collected from clots prepared from five to seven different donors, and the experiments were performed on different days. For each donor and treatment, the microtensometry experiments were performed at least in triplicate, and the results from two or three replicates for each donor were averaged. To establish differences between untreated and treated clots, a paired, one-tailed Student’s *t*-test was performed to calculate *P*-values, for which a value of less than 0.05 was deemed statistically significant.

## 3. Results

### 3.1. Mechanical characterization of blood clots

#### Response of blood clots to uniaxial strain

To measure the stress-strain response of blood clots in the ranges of micronewton force and submicron displacement, we designed a microtensometer that measures tension as the clots are stretched (**Fig. 1A**). The design specifications, engineering details, and calibration of the instrument are presented elsewhere (19). Thrombin-initiated whole blood clots were loaded onto the microtensometer, and secured using magnetic clamps (**Fig. 1A**). Clots having 6 mm gauge length were extended by 3 mm (i.e., 50% strain) at 100 µm/s. Force measurements were recorded simultaneously at 30 Hz. The force-displacement curves were converted to true stress vs. strain plots using cross-sectional area measured in two orthogonal directions. As expected for many biological tissues and fibrous matrix materials, the stress-strain plot for untreated whole blood clots exhibits strain-stiffening behaviour over the tested range of strains (**Fig. 1B**). Next, we evaluated the contribution of platelets to the mechanical behaviour of clots using blood reconstituted with platelet-poor plasma (PPP), and obtained the stress-strain response as outlined above (**Fig. 1B**). The clots from reconstituted blood without platelets were substantially more compliant than whole blood clots. The strain energy density, which is the potential energy stored in the clot due to the work done to deform the clot, is computed as the area under the stress-strain curve. At 50% strain, the strain energy density of whole blood clots was nearly two-fold higher than that of clots formed without platelets, suggesting that the platelets contribute nearly two-fold more work in deforming the clots (**Fig. 1C**). The equivalent linear stiffness of the clot at 50% strain, which is the slope of an ideally linear stress-strain curve having the same strain energy density, was estimated to be 4.4 kPa, which is in the same range of Young’s modulus values reported recently for retracted clots using micropillars (9). The equivalent linear stiffness of clots without platelets was substantially lower at 2.5 kPa.

**Figure 1.**
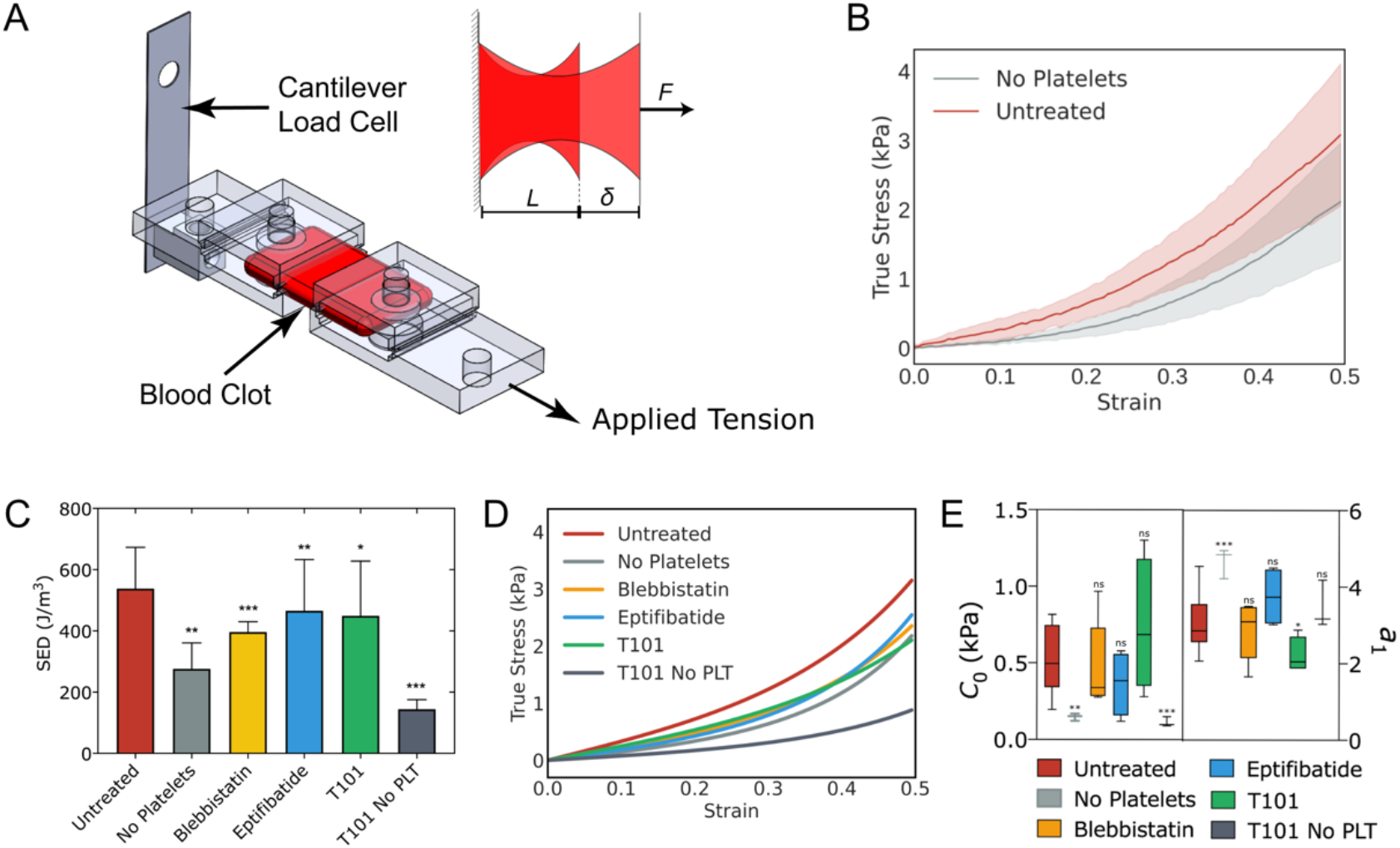
Response of blood clots to uniaxial strain. (A) Oblique view and close-up top-view of clot in between microtensometer grips. One grip is attached to a fixed load cell and the opposite the grip is moved by a nanopositioner.(B) Stress-strain response of clots formed by the addition of thrombin to whole blood or clots formed from platelet poor plasma reconstituted with red blood cells; (C) Strain energy densities at 50% strain; (D) Stress-strain curves fit to the nonlinear exponential model σ = 2*C*_0_*a*_1_exp(*a*_1_ε^2^), where *σ* is stress, *ε* is strain, and C_0_ and a_1_ are model parameters; (E) *C*_0_ and *a*_1_ for untreated and treated clots. Mean ± SD, * *P* < 0.05; ** *P* < 0.01; *** *P* < 0.001.

To parameterize the stress-strain response, we applied an empirical nonlinear fit, using a uniaxial simplification of the Fung strain energy model that has been widely used for soft tissues composed of fibrous networks (20). For this model, the strain energy is given by *u = C*_0_ exp*(a*_1_*ε* ^2^) where *u* is the strain energy density, *ε* is uniaxial strain, and *C*_0_ and *a*_1_ are fitted constants. Differentiating strain energy with respect to strain, we obtain an expression for tensile stress as *σ* = 2*C*_0_ *a*_1_ *ε* exp(*a*_1_ *ε*^2^). The stress-strain experimental data were fit to this equation (**Fig. 1D**). The variables *C*_0_ and *a*_1_ parameterize stiffness at lower strains and strain-stiffening behaviour, respectively. The estimates for *C*_0_ and *a*_1_ for whole blood clots are, as expected, significantly lower than those reported for aortic tissues (21,22) (**Fig. 1E)**. In the absence of platelets, the stress-strain curve fits yielded estimates of *C*_0_ and *a*_1_ that are three-fold lower and two-fold higher, respectively, than untreated clots. Since the stress-strain relationship is nonlinear, the elastic modulus of the clot increases with tensile strain, and can be estimated from the equivalent area under the stress-strain curve. As can be seen from **Fig. S1A**, platelets contribute to nearly two-fold increase in stiffness of the clots over the entire range of strain tested in this work.

#### Effect of platelet contraction and aggregation on clot response to uniaxial strain

Having established the role of platelets to the stress-strain response of blood clots, we sought to determine the role of two key processes, i.e., platelet compaction and aggregation, which are known to impact the overall contraction of a blood clot. To this end, we evaluated the stress-strain response of clots formed with platelets treated with blebbistatin, where clot formation was initiated by the addition of thrombin. Blebbistatin inhibits the binding of myosin II to actin, which is critical to platelet contraction (23). As shown in **Fig. 1C**, we observed that blebbistatin treatment significantly changed the response of clots to uniaxial strain compared to untreated clots. From these plots, we computed the total strain energy density, the nonlinear fit parameters, and the stiffness at 50% strain. The strain energy density was 25% lower in clots treated with blebbistatin compared to untreated clots (**Fig. 1D**).

To evaluate the effect of platelet aggregation on the tensile response of the clots, we similarly tested clots prepared using platelets exposed to eptifibatide, an inhibitor of platelet GP IIb/IIIa, and hence aggregation (24). The uniaxial stress response shows that these clots are weaker than untreated clots, although only modestly so (**Fig. 1C**). The strain energy density for eptifibatide-treated clots was 20% lower than for that of untreated clots (**Fig. 1D**). The parameter *C*_0_ was lower for blebbistatin and eptifibatide treated clots, indicative of overall lower stiffness. The parameter *a*_1_ was only modestly different from that of whole blood clots, indicative of similar strain-stiffening behaviour (**Fig. 1E, Fig. S1A**).

#### Contribution of fibrin-cross linking on clot response to uniaxial strain

To evaluate the effect of fibrin crosslinking, we evaluated the stress response of clots that were polymerized with thrombin but in the presence of T101. T101 is inhibits transglutaminase activity of FXIII, thereby eliminating the lateral crosslinking of fibrin strands (25,26). The strain energy density of non-crosslinked clots was ∼25% lower than that of crosslinked clots. We observed that non-crosslinked clots show significantly lower stress at the same strain, i.e., weaker, compared to the clots formed by the direct addition of thrombin (**Fig. 1D**). Correspondingly, the stiffness of 2.1 kPa at 50% strain was comparable to clots without platelets, indicating that the platelets are as influential as crosslinking in clots initiated with exogenous thrombin. The *C*_0_ value was comparable to untreated whole blood clots but *a*_1_ was lower, revealing the relative importance of crosslinks on stiffness at higher strains and is evident from weaker dependence of elastic modulus with applied strain (**Fig. S1B**). The combined effect of crosslinking and platelets on clot stiffness can be seen from T101-treated blood without platelets. These clots were the weakest with strain energy density three-fold lower than whole blood clots (**Fig. 1C**).

### 3.2. Microstructural characterization of platelet-fibrin interactions in clots

As seen from the mechanical data, platelets contribute to the stiffness of the clots through platelet contraction and aggregation. To understand the contribution of platelets, we analysed clot microstructure through high-magnification fluorescence imaging, with emphasis on platelet-fibrin interactions in clots prepared from platelets that were untreated or treated with blebbistatin or eptifibatide (**Figure 2A**). As evidenced by the intensity of red and green fluorescence due to fibrin and platelets, respectively, interfering with platelet actomyosin or aggregation altered the size of the aggregates and the amount of fibrin present. Interestingly, T101 treatment did not seem to alter either platelet aggregates or the amount of fibrin present on the platelets.

**Figure 2.**
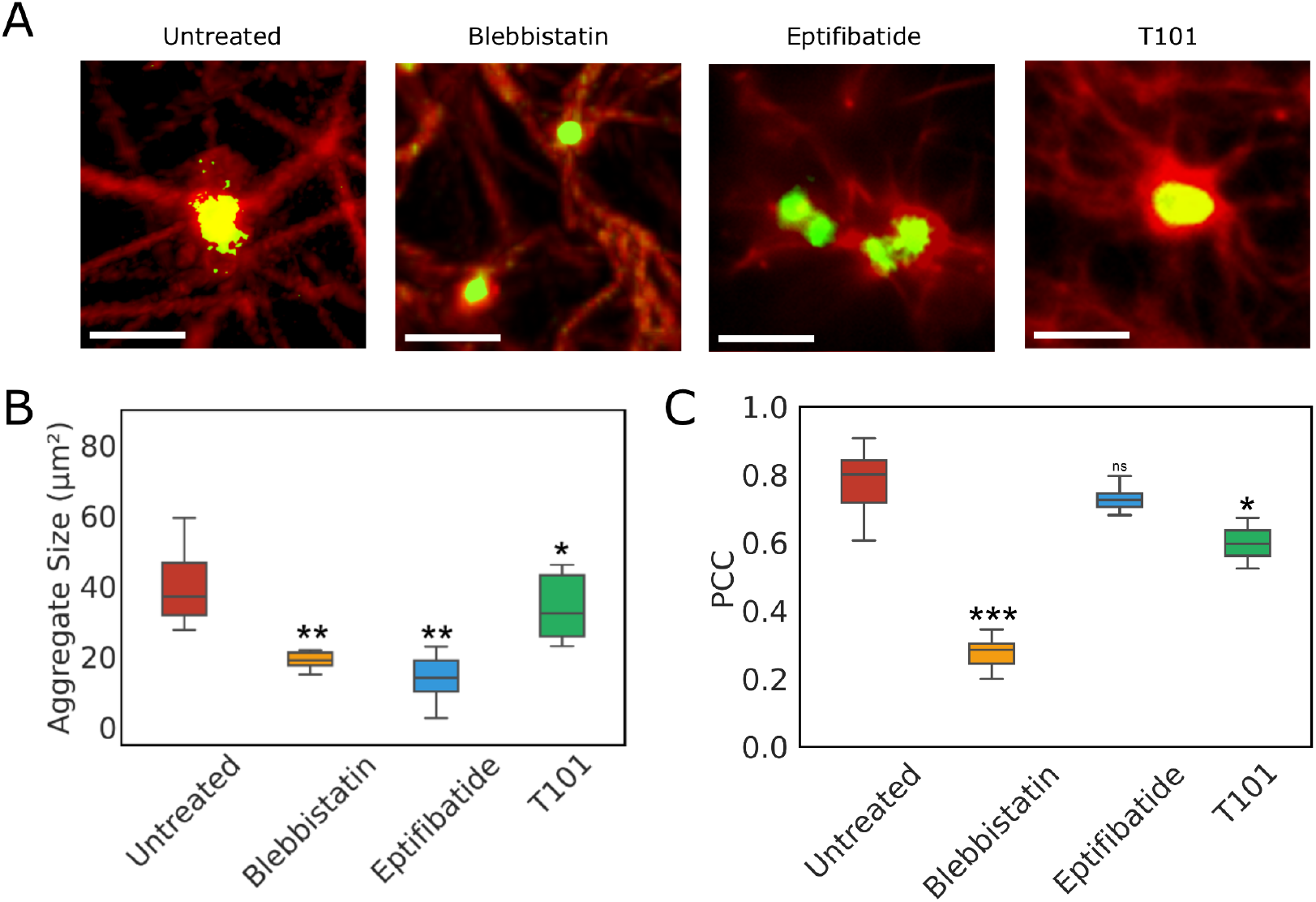
Microstructural characterization of platelet-fibrin aggregates. (A) Clots prepared from platelets that were either untreated or treated with blebbistatin, eptifibatide, or T101. Scale bar = 10 µm; Platelets are labelled with AF488-CD42b (green) and fibrinogen with AF594-fibrinogen (red); (B) Average size of platelet aggregates; (C) Pearson correlation coefficient for the colocalization of platelets and fibrin. Mean ± SD, \**P* < 0.05; \*\**P* < 0.01; \*\*\**P* < 0.001.

To quantify platelet-fibrin interactions, we first estimated the size of platelet aggregates and the amount of fibrin in these aggregates. The average sizes computed based on these images revealed a mean aggregate size of 40 μm^2^ in clots without any treatment, while blebbistatin and eptifibatide treatments reduced the average aggregate size by two-fold and three-fold, respectively (**Fig. 2B**). In contrast, T101 treatment only had a modest effect on aggregate size. Next, we correlated the spatial distributions of the densities of platelets and fibrin based on the overlap in the fluorescence intensities between platelets (green, CD42b-AF 488), and fibrin (red, fibrinogen-AF 596) using the Pearson correlation coefficient (PCC) (**Fig. 2C**). A PCC closer to 1.0 signifies a higher incidence of localization between the green and red fluorophores (27). We observed that in untreated blood clots, 80% of the projected area of the platelet aggregates was also fibrin-rich. In contrast, the spatial overlap between fibrin and platelet densities were low in clots formed from blebbistatin-treated platelets. PCC was only modestly affected by eptifibatide-treated clots, indicating that single platelets or microaggregates co-located strongly with fibrin. A similarly small difference in PCC compared to untreated clots was observed in clots treated with T101, suggesting that fibrin crosslinking may only modestly affect the colocalization of fibrin fibres on platelet aggregates.

### 3.3. Both platelet aggregation and fibrin compaction determine clot stiffness

Upon observing the effect of platelet aggregation and platelet actomyosin on clot stiffness from microtensometry, and also the effect on platelet-fibrin interactions from microstructure studies, we sought to establish a relationship between the two. We estimated the equivalent modulus based on strain energy density at 50% strain. We surmised that the size of the platelet aggregates, and the amount of fibrin bound to platelet aggregates, i.e., PCC, would be critical to clot stiffness. Although neither the size nor PCC individually show a meaningful correlation with equivalent modulus, we discovered that the product of these quantities, which estimates the density of fibrin bound to platelet aggregates, is tightly correlated with the equivalent modulus of the clot (**Fig. 3**). The effect of fibrin bound to the platelet aggregates on elastic modulus was independent of applied strain (i.e., slope of 52 to 57 kPa/μm^2^ over 10% to 50% strain) indicating that the fibrin-platelet interactions determine the effective mechanical behaviour of the clot. These data show that both platelet aggregation and actomyosin contraction result in comparable densities of fibrin-bound platelet aggregates, and they contribute similarly to increasing the stiffness beyond the baseline no-platelet clot, albeit by different mechanisms. Interestingly, disrupting fibrin crosslinking with T101 had two effects: it not only reduced the amount of fibrin bound to the platelet aggregates but also lowered the equivalent modulus from what may be expected of fully crosslinked clots.

**Figure 3.**
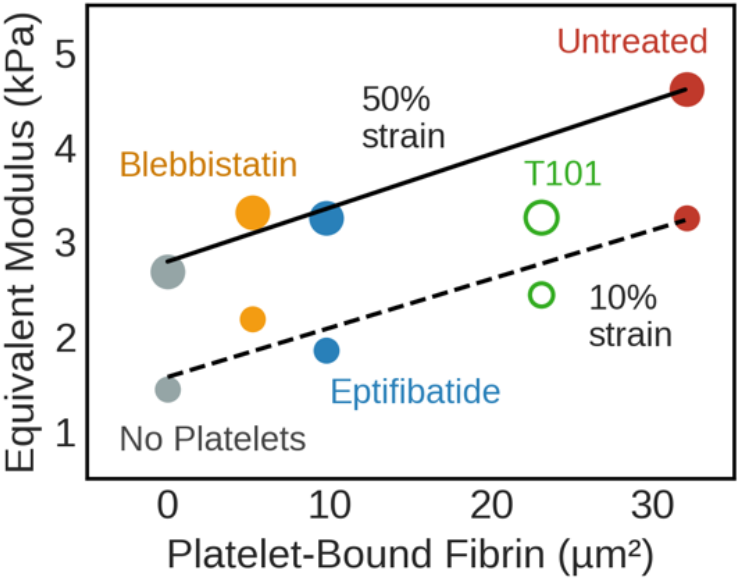
Platelet-fibrin interactions determine clot stiffness. (A) Elastic modulus correlates linearly with the density of platelet-bound fibrin at various applied strains. The lines represent least-squares best fits without T101 (with *R*^2^ = 0.90 at 10% strain and *R*^2^ = 0.97 at 50% strain, respectively).

### 3.4. A computational framework for the contribution of fibrin-bound platelet aggregates on clot stiffness

#### The discrete prestressed fibre model

Having established a correlation between the density of fibrin-bound platelet aggregates and strain energy density, we developed a computational model to interrogate the underlying physical basis of this relationship. A discrete prestressed fibre model and its functional components that include a fibrin-bound platelet aggregate are shown in **Fig. 4A**. The platelet-fibrin aggregate is represented by a contractile element with an elastic modulus *E*_*c*_, which may be different from the modulus of elastic modulus *E*_*f*_ of distal fibres. The ratio between moduli is defined as relative modulus, *β* = *E*_*c*_/*E*_*f*_. The fibres are bound to platelet GP IIb/IIIa receptor clusters at the focal adhesion sites located at a radial distance *R*_0_ from the centre of the aggregate. Contractile forces during clotting due to actomyosin activity draws fibres inward, and are retracted to a tighter aggregate of radius *R*_*c*_. This contraction induces a prestressed condition in the vicinity of the aggregate, which is retained by the clot even in the absence of an externally applied load. A quasi-static model of platelet contraction is a realistic approximation since the timescale of contraction-relaxation of non-muscle myosin IIA is much smaller than the timescale of matrix contraction (28). This prestress is due to plasticity of the overall cell-matrix interaction, as has been observed for cells encapsulated in collagen and fibrin gels (29). **Fig. 4B**, superimposed on the average stress-strain curve of the clots without platelets (i.e. fibrin network), illustrates how prestress can influence the effective modulus. In the absence of prestress, an externally applied strain *ε*_*ext*_ results in a nominal stress *σ*_*nom*_. The local modulus at the corresponding nominal strain *ε*_*nom*_ is *E*_*nom*_. However, when prestress *σ*_*pre*_ and prestrain *ε*_*pre*_ are present, the addition of the same externally applied strain results in a higher effective stress *σ*_*eff*_, and the local effective modulus *E*_*eff*_ at the (total) effective strain *ε*_*eff*_ is steeper. The corresponding strain energy density, computed as the area under the curve up to the effective strain, is also substantially larger than nominal.

**Figure 4.**
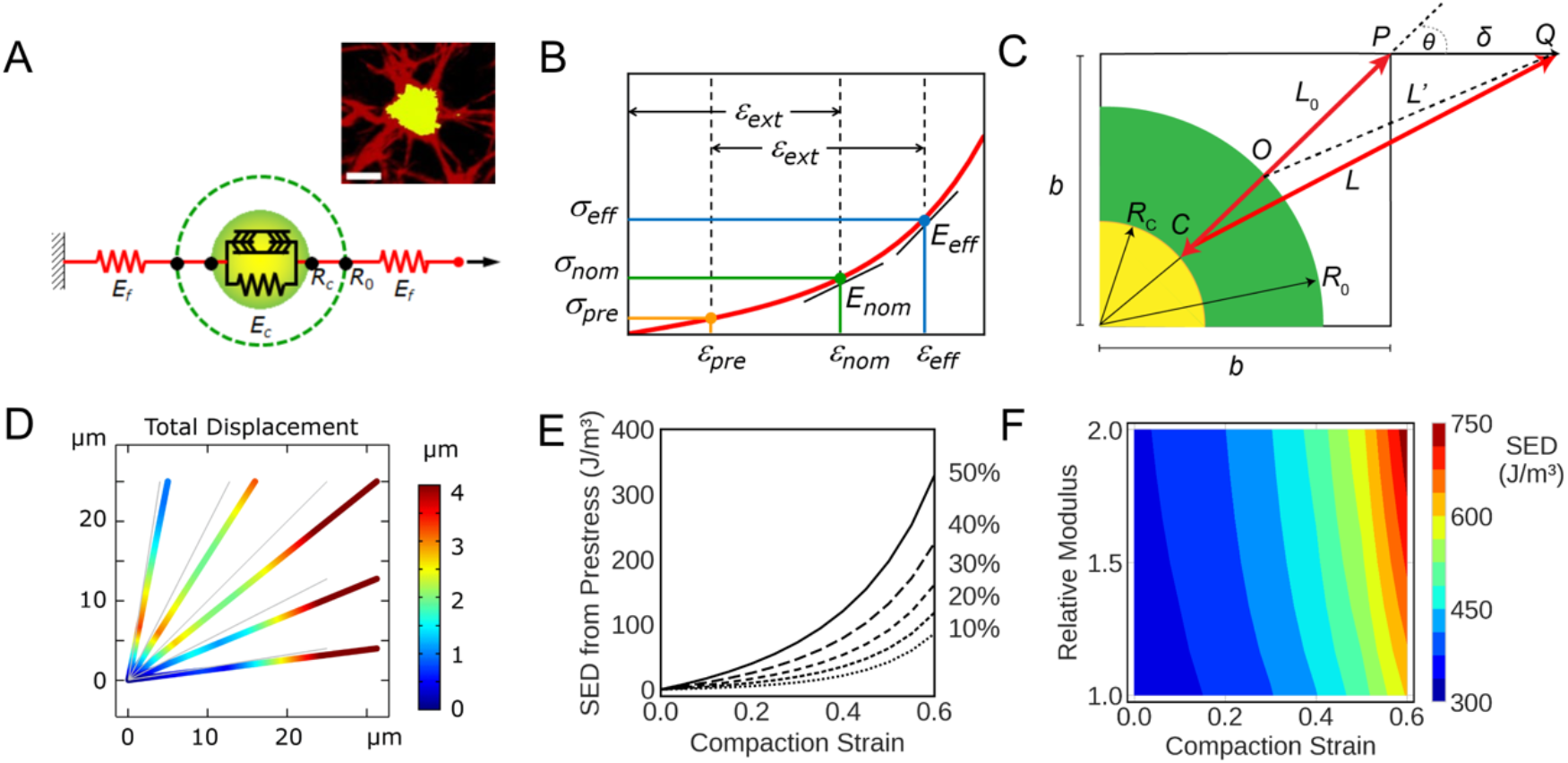
Discrete prestressed fibre model for clot subjected to uniaxial strain. (A) The actomyosin contractile elements in platelets compact the fibrin fibres onto the aggregates; Representative image of a single platelet aggregate with fibres emanating from the aggregate (Scale bar: 5 μm) (Inset); (B) Compaction strain is added to externally applied strain, such that the effective modulus *E*_*eff*_ is higher than nominal.; Euclidean geometric representation of a fibrin fibre compacted by platelet contraction; (D) Representative finite element simulation of fibre deformation in the vicinity of a platelet aggregate, shown with 5 µm aggregate radius, 0.5 compaction ratio, 2.0 relative modulus, and 25% applied strain; (E) Strain energy density due to prestress as a function of compaction strain for applied strain up to 50%; (F) Predicted strain energy density as a function of compaction strain and relative modulus.

The interaction between a representative fibrin fibre bundle and a platelet aggregate is depicted in **Fig. 4C**. Platelet GP IIb/IIIa-fibrin(ogen) interactions anchor the fibrin fibres to the platelet surface. The fibre of an initial length *L*_0_ is bound to the platelet aggregate at a focal adhesion at point *O*. A quarter-circle that represents a single platelet aggregate with two orthogonal planes of symmetry is shown within a corresponding quadrant of a square having side length *b*. The fibre is oriented at an arbitrary angle *θ* with respect to the direction of applied external strain. The fibre *OP* experiences the combined effects of prestrain and applied external strain, resulting in a geometrically nonlinear response to strain. The individual fibres are modelled as linear elastic materials obeying Hooke’s law.

The global stiffness matrix for two fibres in series is written as shown in equation (1), where node 0 is at the centre of a platelet aggregate, node 1 is at the focal adhesion, and node 2 is the free end subject to external strain. *F* represents nodal forces and *d* represents nodal displacements along the axial direction *x*. Spring constants *k*_1_ and *k*_2_ are associated with the platelet aggregate and (unbound) fibrin fibre, respectively.

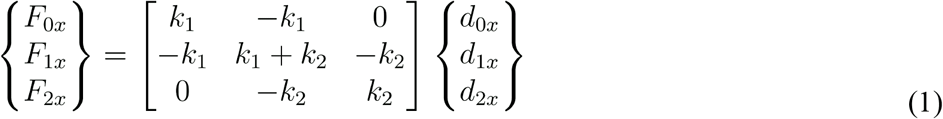

In order to represent prestress, we add a contractile force *F* = −*αEA*, and the focal adhesion (i.e., node 1), where the elastic modulus *E* and cross-sectional area *A* pertain to the platelet aggregate. The parameter *α* is the compaction strain, defined as the extent of aggregate compaction from an initial radius *R*_0_ to a compacted radius *R*_*c*_ (i.e., *α* = (*R*_0_ − *R*_c_)/*R*_0_). At node 2 (i.e., the location where external strain is applied), the fibre is subjected to a horizontal displacement *δ*. Applying these assigned conditions, the equilibrium solution to axial force along the fibre assembly is as shown in equation (2), with details provided in the Supplementary Information.

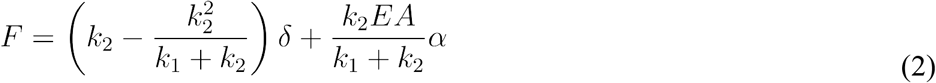

In the presence of prestrain due to compaction (*α* > 0), the second term in the force solution above predicts a higher axial force along the fibre with increasing compaction strain. From this solution, the corresponding strain energy density can be calculated for any combination of relative modulus *β*, compaction strain *α*, and applied uniaxial strain *ε* = *δ/b*. The multi-directional orientation of fibres was accounted for by using coordinate transformations for fibres that are oriented off-axis from the direction of externally applied strain, as explained in the Supplementary Information.

#### Effect of fibrin compaction and fibre modulus on the strain energy density of the clots

Finite element simulations were performed using truss-element fibres at evenly spaced angles in a quadrant with two-plane symmetry. The model showed convergence in strain energy density with as few as three fibres per quadrant, and simulations presented here used five fibres per quadrant (**Fig. S3A**). **Fig. 4D** shows the distribution of local deformation of the fibrin fibres, under the combined effect of platelet contractile strain and externally applied strain. As expected, the fibres oriented in the direction of externally applied strain experienced the largest magnitudes of local displacement. For the same applied strain, the absence of platelets resulted in smaller local deformation of the fibres when compared to the deformation in the presence of platelet contraction (**Fig. 4D** vs. **Fig. S3B**). In the absence of platelets, the strain energy density of the fibrin fibres is purely due to externally applied strain, and scales approximately as the square of strain (**Fig. S3C**).

We computed strain energy densities using the five-element model for a range of compaction strains and externally applied strains. The compacted radius *R*_*c*_ and the bounding-box size *b* for each clot type were set accordingly to account for the average space occupied by a single platelet aggregate, based on the average density of aggregates measured from image analysis. Since strain energy density is additive, the effect of applied strain can be subtracted from the total strain energy density to obtain the effect of compaction strain. As shown in **Fig. 4E**, we observed that the contribution to strain energy density from compaction increases exponentially with an increase in compaction strain, and is more pronounced at higher externally applied strains because the fibres are effectively subjected to strains in a steeper region of the stress-strain curve.

The above simulations were performed assuming that the elastic modulus of the fibrin fibres is twice as much upon compaction on the platelet surface as when they are not bound to platelets. Since the actomyosin stress fibres are indeed rigidity sensors, and respond to increase in local stiffness by exerting larger traction forces, we also considered the effect of an increase in the elastic modulus of fibrin fibres that get compacted on the platelet surface due to contraction (30) (**Fig. 4F**). We noticed that the total strain energy density was only modestly affected by the platelet-bound fibrin as being stiffer than unbound fibrin, over a broad range of compaction strains. From these simulations, we demonstrated that our discrete fibre model accounts for the effects of both compaction strain due to platelets and strain due to externally applied tension, in agreement with the total strain energy density measured experimentally. It should be noted that modelling fibrin adsorbed on platelet aggregates as merely passive spring elements would have failed to capture experimental observations (**Fig. S2**). In the absence of compaction strain, even a ten-fold increase in elastic modulus of fibrin bound to the platelet aggregates produced a barely perceptible increase in strain energy density.

#### Fibrin compaction on platelet surface due to aggregation and actomyosin contraction increases the strain energy density of clots

Next, we sought to obtain estimates of compaction strain by matching the total strain energy density from model predictions against experimentally measured values. Since the simulations show that the relative modulus has only a modest effect on strain energy density over the experimental range (i.e., 300 J/m^3^ to 750 J/m^3^), without loss of generality, we considered a model wherein the modulus of the fibre increased by two-fold upon compaction (**Fig. 4E**). To compare the model predictions to experimental stress-strain data, we superimposed the experimentally determined strain energy densities at different applied strains (computed from **Fig. 2A**) on to the model predictions (**Fig. 5A)**. As the applied tensile strain increases beyond approximately 20%, and the resistance to deformation is due to fibrin fibres bound to platelets, the intersections between experimental and predicted strain energy densities converge at a characteristic compaction strain of approximately 0.6.

**Figure 5.**
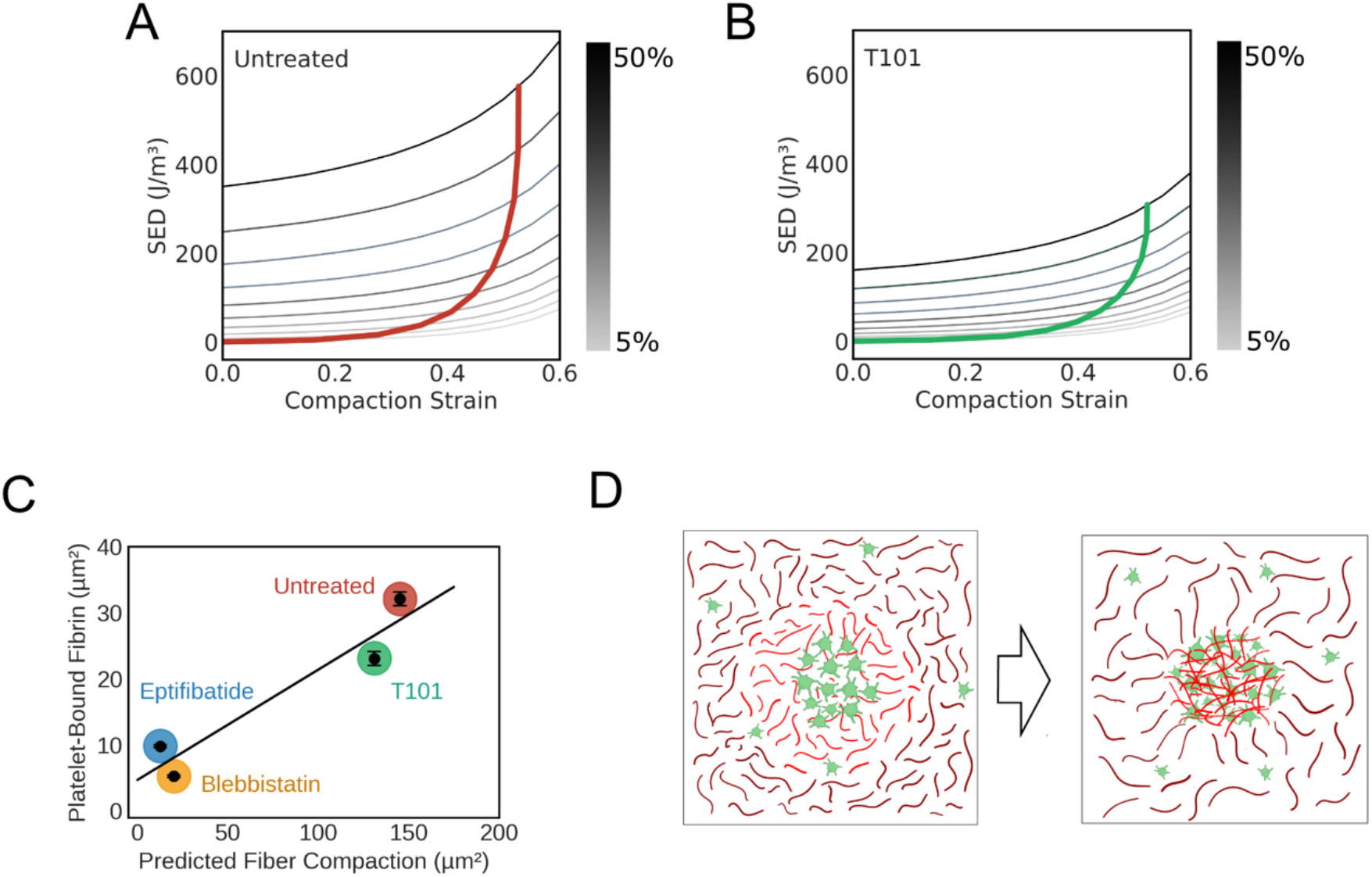
Discrete prestressed fibre model predicts the amount of fibrin fibre compacted by contracting platelets. (A, B) Mapping of experimentally determined strain energy density onto model predictions for crosslinked (A) and uncrosslinked clots (B); (C) Correlation between experimentally determined fibrin density on platelet surface and predicted fibre compaction based on area reduction (*R*^2^ = 0.91); (D)Schematic representation of predicted fibre compaction by platelets. Platelet aggregates sequester fibrin fibres into dense nodes, thereby prestressing the fibre network.

Similarly, we estimated the compaction strain for clots formed from platelets treated with blebbistatin or eptifibatide (**Fig. S4A and Fig. S4B**). Both blebbistatin and eptifibatide treatments significantly reduced the compaction strain to nearly three-fold lower than for untreated platelets, indicating that both platelet aggregation and actomyosin contraction are critical in increasing the effective modulus of the clot. As shown in **Fig. 5B** for clots treated with T101, the predicted compaction strain was comparable to that of untreated clots, indicating that disrupting crosslinking of fibrin fibres does not alter the ability of the platelets to contract the fibrin fibres. For computing the compaction strain in T101-treated clots, we used the elastic modulus of fibrin fibres estimated from clots formed from blood depleted of platelets and also treated with T101. The elastic modulus of these clots formed from blood depleted of platelets but treated with T101 were much lower than the elastic modulus of clots treated with T101, indicating the significant impact of fibrin crosslinking on clot properties.

#### Fibrin compaction predicted from the model correlates with proportion of fibrin on platelet aggregates

As discussed above, the compaction strain predicted by the discrete fibre model provides a measure of fibrin densification on the platelet surface. In **Fig. 5C**, we observe that this is indeed the case, as the total amount of fibrin sequestered on to the platelet surface due to compaction strain is linearly correlated to the fibrin-bound to platelet aggregates as estimated by the product of PCC and aggregate area. This observation suggests that the extent of colocalization on aggregates observed by microscopy may serve as a quantifiable metric for estimating fibrin compaction that leads to prestress in the clots.

We note that the amount of fibrin compacted by platelets decreases by five-fold if the actomyosin contraction or aggregation is inhibited, as evident from blebbistatin or eptifibatide treatments respectively. Interestingly, the amount of fibrin compacted by platelets was not as much influenced by the ability of fibrin to crosslink, suggesting that the decrease in elastic modulus of T101-treated clots is caused mainly by lack of crosslinking in the matrix, rather than by the inability of fibrin to interact with platelet aggregates.

## 4. Discussion

Both conventional rheometry and more recent micropost measurements have shown that the presence of platelets increases the stiffness of clots by, depending on the measurement techniques, two-fold to five-fold in the 0.8 kPa to 20 kPa range (9,26,31). The increase in clot stiffness is a result of clot retraction, volume shrinkage, and plasma exudation driven by platelet contraction. As platelets contract, they appear to tug along the fibrin fibres, and actively remodel the fibrin network in their vicinity (10,32).

However, the biophysical mechanisms by which these platelet-fibrin interactions increase clot stiffness is unknown. We show that the work done by contracting platelets in remodelling the fibrin fibres is stored as strain energy density even before an external load is applied, and is a critical contribution to the mechanism by which the clots resist deformation (33). Analogous to macroscopically familiar structures such as a rope hammock or an anchor capstan, our experimental data and modelling support the hypothesis that the discrete prestressed nodes composed of fibrin-bound platelet aggregates are key to the structural stability of the clots (34) (**Fig. 5D**).

The local contractile strain exerted by platelets on the fibrin network is somewhat similar to from the contractile strain exercised on the extracellular matrix by contracting cells which remodels fibres in the immediate vicinity of the cells and increases the overall stiffness of fibrous-matrix gels (35,36). However, there are several important distinctions between the contractile strain exerted by platelets vis-à-vis fibroblasts: First, while cells rearrange the fibrous networks of the extracellular matrix in their immediate vicinity, platelet filopodia draw and spool the fibrin fibres inward so much that the platelets are wrapped in fibrin fibres, as evidenced by the colocalization indices. We have deliberately used the term ‘compaction strain’ instead of ‘contractile strain’ to underscore this distinction. Second, compared to cellular traction forces, the contractile forces generated by single platelets are more than an order of magnitude lower in the 2 nN to 20 nN range, suggesting that platelet aggregation may be key to compensating for the smaller size and force (37–39) compared to fibroblasts. Our experimental data suggest that this is indeed the case: (i) although disrupting platelet aggregation using eptifibatide did not alter platelet-fibrin interactions (i.e., PCC) at the single platelet level, it significantly weakened the clot; (ii) based on the increase in the stiffness of whole blood clots (4.4 kPa) compared to no-platelet clots (2.5 kPa), and using an average platelet aggregate diameter of 7 μm, we can estimate the mean contractile force exerted by a platelet aggregate to be in the few hundreds of nanonewtons, which is in a range comparable to cellular traction forces (40). Third, simultaneous and cooperative compaction of fibrin by platelet aggregates is likely to be important in increasing clot stiffness, since neither activated but poorly aggregating platelets nor a priori aggregated platelets retract efficiently (41,42). Our computational results confirm that this is indeed the case since only the model accounting for active platelet contraction, but not passive stiffening of platelet-fibrin aggregates, is capable of matching experimental strain energy density measurements.

A prominent result from this study is that fibrin that is taken in by platelet aggregates from their immediate vicinity is sufficient to increase resistance to deformation substantially for the entire clot, even though most of the fibrin network may remain agnostic to platelet contraction. As a corollary, if the platelet-fibrin aggregates were not prestressed, to provide the strain energy density equivalent to that of a whole blood clot, the entire fibrin network would have to be twice as stiff. This increase in stiffness would have to arise purely from fibrin polymerization and crosslinking, which is a physiologically undesirable process. In light of growing interest and relevance on the spatial heterogeneity in the composition of *in vivo* clots to their strength (43,44), our results present some of the first insights into the origins of clot response to external forces. We expect that these insights will lead to more detailed expositions of clot deformation through the use of appropriate image-based 3D models of the network accounting for non-linear fibre properties as well as platelet-fibrin interactions.

In summary, we have measured the elastic moduli of blood clots subjected to uniaxial tensile strain using microtensometry, and we have estimated the contribution of platelet contraction and aggregation to clot stiffness using specific inhibitors of non-muscle myosin IIA, aggregation, and crosslinking. Our results show that the platelet contribution to clot stiffness is dominated by nodes in the fibrous matrix that are composed of platelet aggregates and compacted fibrin. The concept of nodal prestress introduced in this work provides insights into how fibrin sequestering during clot retraction serves as a key stabilizing factor in clot mechanics. The experimental and computational framework presented in this work, for the first time to our knowledge, not only measures tensile properties of clots but also provides a simple yet powerful model to establish mechanobiological relationships between intracellular stress and tissue-scale external loads.

## Acknowledgments

Assistance with data collection from C. Puligundla, L.A. Resnikowski, J. Alzaghari, and S-M. Saw is gratefully acknowledged.

## Author contributions

SJP and WE performed experiments and analysed data. SJL performed computational studies. SJL and AKR conceived the research, designed experiments, performed data analysis, procured funding, and wrote the manuscript. All authors approved the manuscript.

## Supplemental Text

### S1. Materials and Methods

#### Acquisition of human blood

Blood was obtained from donor volunteers according to the protocol approved by the Institutional Review Board (F16134) at San José State University. The donors were 18 to 25 years of age, healthy, free of any known bleeding or cardiovascular disorders, nondiabetic, and not under any platelet-function altering medication. The blood was drawn in 3.2% buffered sodium citrate vacutainer tubes (BD Biosciences, San Jose, USA).

#### Preparation of blood clots

500 µL of blood was recalcified by the addition of sterile-filtered CaCl_2_ (Millipore Sigma, St. Louis, MO, USA) at pH 7.4 to a final concentration of 20 mM, and restriction grade thrombin (Millipore Sigma, St. Louis, MO, USA) to a final concentration of 0.2 U/mL in wells of a 25 mm x 75 mm chamber slide (Nunc, Rochester, USA) and allowed to clot for 90 min at room temperature. For designated experiments, the following inhibitors were added to blood, and incubated for 30 min at room temperature with gentle rocking (Vari-Mix Platform Rocker, Thermo Fisher Scientific, Waltham, MA, USA) at approximately 2 Hz before recalcification: blebbistatin (Calbiochem, San Diego, CA, USA) at 300 μM; eptifibatide (Bio-Techne Corp., Minneapolis, MN) at 10 µM; or T101 (Zedira GmbH, Germany) at 10 µM. To investigate the effect of platelets in clotting, red blood cells (RBCs) were separated by centrifugation for 15 min without brakes at 250 RCF, 7 rad/s^2^ acceleration (Eppendorf). After separating the RBCs from platelet rich plasma (PRP), platelet poor plasma (PPP) was obtained by further centrifugation of PRP for 20 min without brakes at 1500 RCF, 7 rad/s^2^ acceleration. The RBC was added back to PPP to restore the haematocrit to original levels in the whole blood.

#### Measurement of clot response to uniaxial strain

##### Design and fabrication of custom microtensometer device

The microtensometer consists of a motorized nanopositioner (MP-285, Sutter Instrument, Novato, CA, USA), a full-bridge thin cantilever load cell (113 g, LCL Series, OMEGA Engineering, Norwalk, CT, USA), and a signal conditioner DI-1000U (Loadstar Sensors, Fremont, CA). The nanopositioner has a 25 mm range resolution of 40 nm per step. Speeds are selectable from 20 µm/s to 2.9 mm/s. The DI-1000U has 24-bit resolution and up to 80 Hz sampling rate.

An aluminium clamp with a recessed crevice that allowed for two points of contact was found to provide consistent clamping. A magnet was press-fit inside the lower jaw of each clamp to create a clamping force with the upper jaw that had a steel screw attached. The magnetic clamping force was adjustable to prevent tearing while ensuring no slip, by adjusting the height of the screw and locking it in place with a counter-tightened nut.

##### Acquisition of force-displacement data

Clots were loaded in between two clamps with fine adjustment enabled by manually operated 3-axis micromanipulators (Cascade Microtech, Beaverton, OR, USA). Each clot began at 6 mm gauge length, and load cells were tared to zero before applying extension. The force recording from the data acquisition software (SensorVUE, Loadstar Sensors, Fremont, CA, USA) and actuation of the nanopositioner were started simultaneously and force measurements were recorded at 30 Hz. The clots were continuously strained to 3 mm at 100 µm/s which was 50% of the initial 6 mm gauge length. Each single experiment was completed within 3 minutes for any given clot, and replicate clots (typically three) in each set of runs were kept under plastic cover to prevent drying.

##### Estimation of cross-sectional area of the clot

Prior to applying strain, one digital microscope camera was set to capture top-view images of the clot to extract top cross-sectional dimensions, and a second digital camera was used to capture side-view images to extract thickness. The cameras captured images during the strain loading, each 8 s apart, to obtain true stress from force measurements. The cross-sectional area from the sequence of images were used to convert force measurements to true stress values. ImageJ software (NIH) was used to measure the three top-view images for the width *w* and two side images for thickness *t*. The area of the clot at 10 intervals along the length was calculated by measuring the width and thickness from the top and side view images taken.

##### Structural characterization of clots Fluorescence microscopy

To visualize clot microstructure, platelets and fibrinogen were labelled with AF488 and AF596 fluorophores, and 100 μL of blood was used to form clots as described above. Platelets were incubated with AF488-conjugated CD42b antibody for 20 min prior to the start of clotting, and blood was doped with 5% AF596 conjugated fibrinogen, resulting in a final platelet concentration of 4 mg/mL. After 1.5 h, the clots were imaged at 63x magnification (0.95 NA) using an oil immersion lens (DMi8, Leica Microsystems, Germany). Z-stack images were collected from at least five different locations for each clot.

##### Image analysis

The fluorescence images were processed and analysed offline using Image-Pro (Media Cybernetics, Rockville, MD, USA) or ImageJ (NIH). After subtracting the background noise by applying a Gaussian filter, the size of platelet-fibrin aggregates, and the platelet-fibrin colocalization in the aggregates (i.e., Pearson Correlation Coefficient) were estimated.

##### Estimation of strain energy densities from finite element simulations

Strain energy densities at different values were estimated using finite element simulations (COMSOL Multiphysics, Stockholm, Sweden) with parametric sweeps along values of (radial) compaction strain and (uniaxial) applied strain. The model was run in 2-D space with quasi-static deformation. Geometric nonlinearity was enabled such that the direction of tension remained axial as each element changed orientation throughout the prescribed extension. Young’s modulus was assigned distinctly to interior and exterior fibres (with respect to the radial boundary for the platelet aggregate), and Poisson ratio was kept constant at a value of 0.49. The effect of compaction was simulated by prescribing (negative) thermal expansion only for the interior fibres. Equal displacement and force equilibrium are maintained at the node joining each “within aggregate” and “outside of aggregate” fibre. Element nodes for the two-plane symmetry quadrant are shown in **Fig. S2B**. The common node at the origin is fixed and right-side nodes are assigned a prescribed strain corresponding to experimental values. The two nodes along the top side are assigned correspondingly proportional strains according to their horizontal positions (i.e., mimicking a roller condition for a continuous body).

A baseline model was first calibrated using the average stress-strain curve of a no-platelet clot (i.e., zero compaction strain) to determine the appropriate Young’s modulus that resulted in the experimentally known strain energy density at each 10% increment of external uniaxial strain between 10% and 50%. The corresponding Young’s modulus values varied between 3.5 kPa and 4.7 kPa over this range of strain. A similar but separate baseline model with no platelets was also established for clots treated with T101, for which Young’s modulus values varied between 1.8 kPa and 2.1 kPa. For all clots involving platelets (i.e., untreated, blebbistatin, eptifibatide, T101), geometric dimensions were set to match the compacted radius that was determined from image analysis (**Fig. 2**). The relative size of the model quadrant was customized according to the spatial density of aggregates, which was determined from image analysis by computing the average number of aggregates per unit area for each type of clot. The corresponding lengths of fibrin fibres were set to maintain the relative size of a single platelet aggregate, to ensure correct proportion of aggregate area with respect to the bounding-box quadrant of the fibre model. The corresponding relative sizes (i.e., aggregate radius divided by quadrant box edge length) for untreated, blebbistatin-treated, eptifibatide-treated, and T101-treated clots were 0.087, 0.100, 0.103, 0.118, respectively.

### S2. Analytical expression for force on prestressed fibrin fibres due to uniaxial tensile strain

Each fibre element in Fig. 4B is assumed to have a linear relationship between axial stress *σ* and axial strain *ε* according to Hooke’s law *σ* = *Eε*, where *E* is the elastic modulus. Letting *û* represent the axial displacement function along the element in the *ê* direction, the strain in the element is *ε* = *dû/dê*, where the caret symbol (^) signifies a scalar direction that is always aligned with the longitudinal axis of the considered element. Equilibrium requires that the axial force *F = σA* be constant along the element, where *A* is the cross-sectional area of the element. Accordingly, the governing differential equation for the element is as shown in equation S2a.

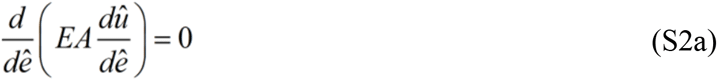

Assigning a linear displacement function for *û* and writing in terms of the nodal displacements 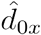 and 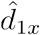 of the end nodes in local coordinates, the displacement function in matrix form is as shown in equation S2b.

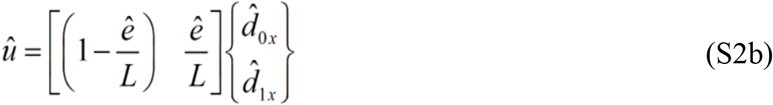

Equation S2c shows the corresponding stiffness equation for the element between node 0 and node 1, in terms of nodal forces 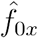 and 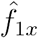.

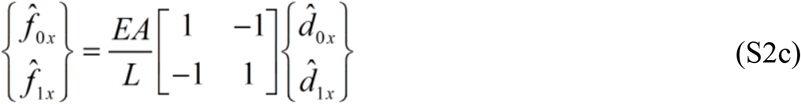

Similarly, equation S2d shows the notation for a second element between node 1 and node 2.

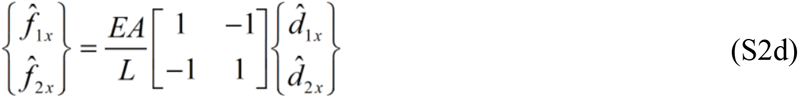

By superposition and using distinct subscripts 1 and 2 for the respective values of area, modulus, and length of each fibre, the global stiffness equation for the two elements in series is shown in equation (S2e).

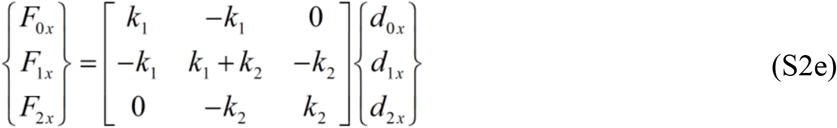

By symmetry of placing node 0 at the origin of the quadrant, *d*_0x_ = 0. For tensometric loading, the end displacement *d*_2x_ is a prescribed value *δ*. There is no externally applied force at node 1, such that *F*_1x_ = 0. With these boundary conditions, the solution reduces to two equations and two unknown (S2f).

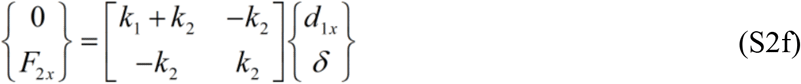

The solution to the scenario with only a prescribed external displacement is straightforward with *d*_1x_ = *k*_2_*δ*/(*k*_1_+*k*_2_) and the axial force *F*_2x_ = *F* as shown in equation (S2g).

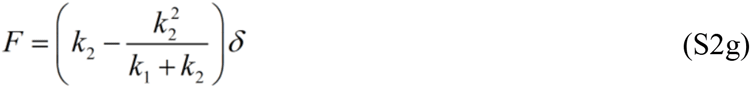

The strain energy in each specific element *i* can be computed from stored elastic energy ½*k*_i_*u*_i_^2^ nodal displacements, where the axial displacement *u*_*i*_ is computed from the difference in nodal displacements (e.g., *u*_1_ = *d*_1x_ -*d*_0x_). The corresponding strain energy density for each element or for the overall system is computed by dividing by respective volume.

We make two important modifications to the basic two-element solution above for our prestrained discrete fibre model. First, at node 1 (i.e., location of the focal adhesion) we add a contractile force *F = -αEA*, for which the elastic modulus *E* and cross-sectional area *A* pertain to the platelet aggregate. Entering this force into the global stiffness equation and simplifying reduces the system to two equations (S2h).

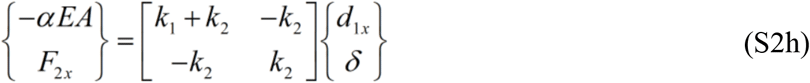

The extent of aggregate compaction from an initial radius *R*_0_ to a compacted radius *R*_c_can be described either by the compaction strain *α* or alternatively in terms of compaction ratio *ζ* = *R*_0_/*R*_*c*_ = 1/(1-*α*). With compaction, the displacement at node 1 becomes *d*_1x_ = (*k*_2_*δ* -*αEA*)/(*k*_1_ + *k*_2_). Equation (S2i) shows the modified solution for axial force when prestrain is taken into account.

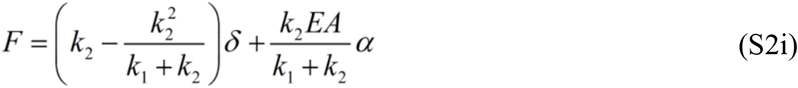

Second, we account for multi-directional orientation of fibres using coordinate transformations for fibres that are oriented off-axis from the direction of externally applied strain. Coordinate transformation is achieved by using a rotation matrix to allow any element oriented at an angle *θ* (measured counter clockwise from the global x-axis) to be mapped to global displacements *d*_*x*_ and *d*_*y*_ as shown in equation S2j.

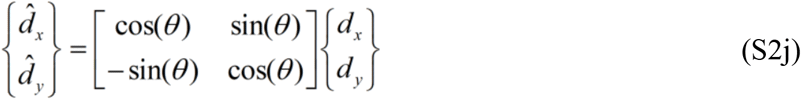

This prestress model formulation was verified using finite element analysis (FEA) by taking advantage of the “thermal force” analogy used to model thermal expansion and contraction. The FEA computational solutions matched analytical solutions exactly for the entire range of relative size, relative modulus, compaction ratio, and orientation angles.

**Figure S1.**
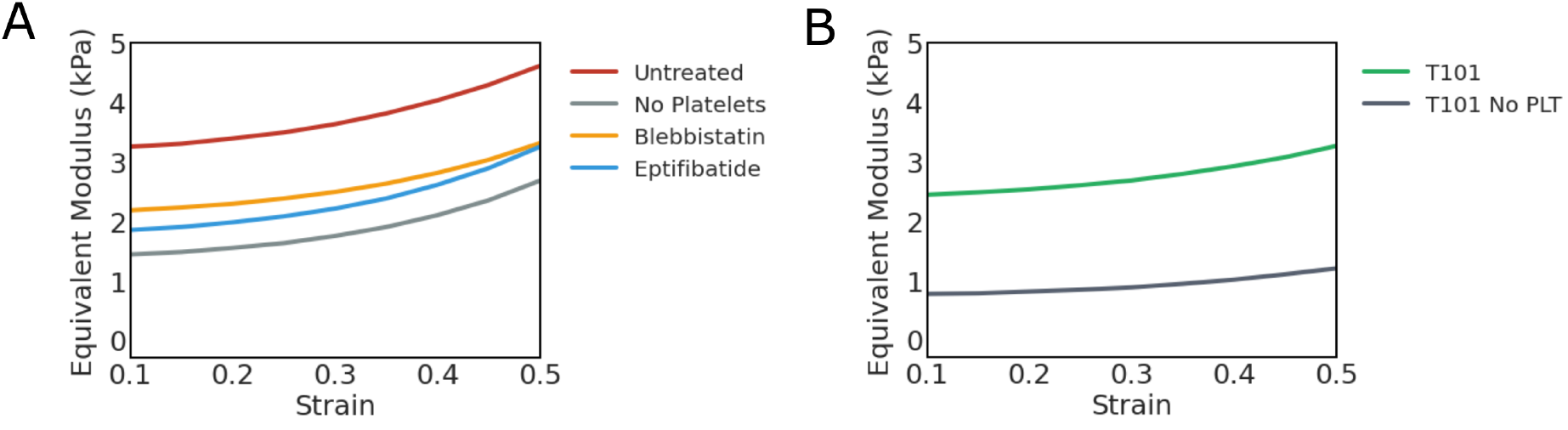
Equivalent modulus of the clots evaluated from the nonlinear exponential model at indicated strains.

**Figure S2.**
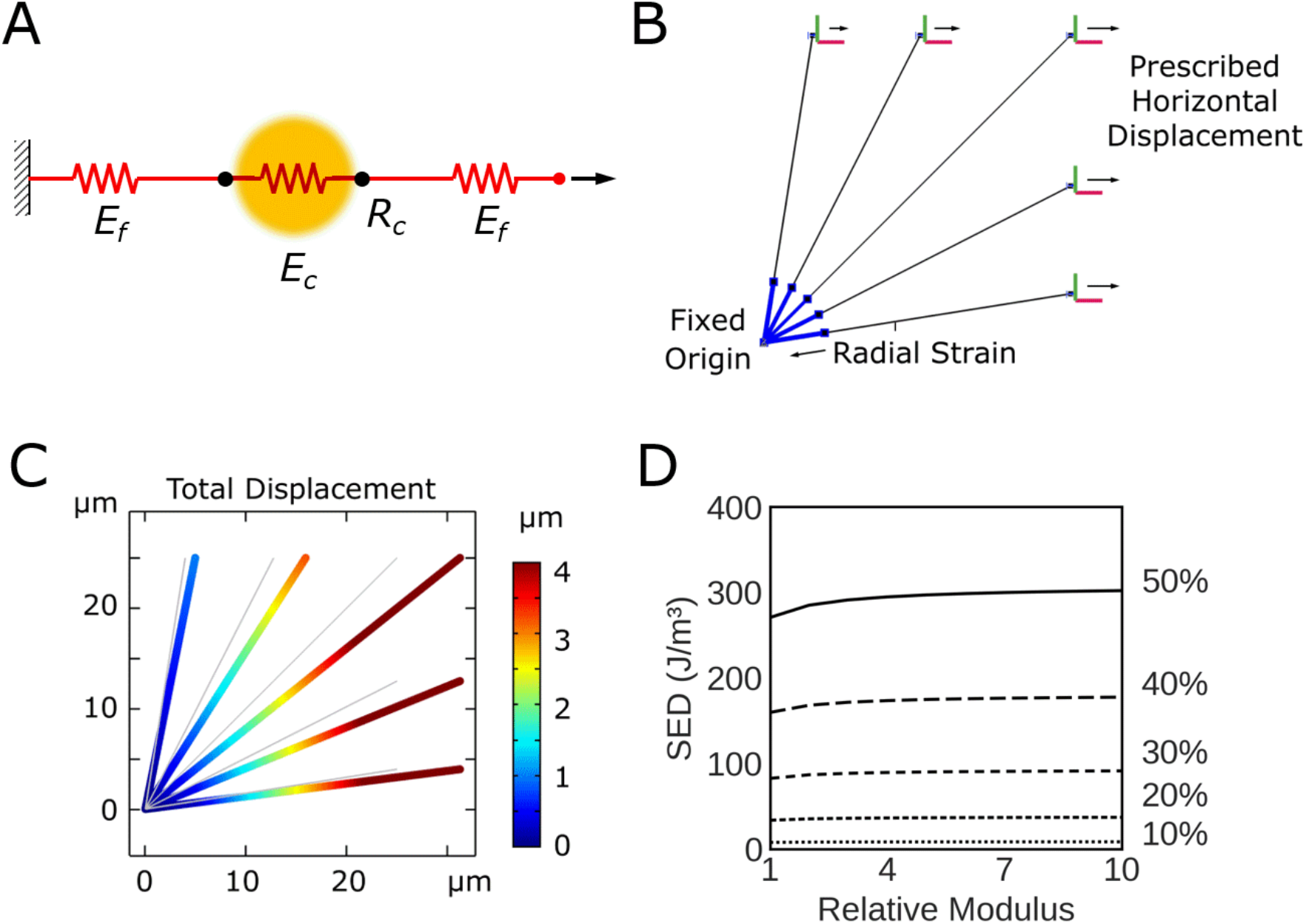
Passive fibre adsorption model. (A) Fibrin fibres (red) are connected to fibrin-bound platelets (yellow), and both fibres are modelled as springs with different stiffnesses. In this passive model platelets do not exercise any active contractile stress; (B) Description of model elements and boundary conditions; (C) Representative deformation of a five-fibre model with 5 µm aggregate radius and relative modulus of 10, pulled to 25% uniaxial strain. (D) Strain energy density as a function of relative modulus, at various applied strains up to 50%.

**Figure S3.**
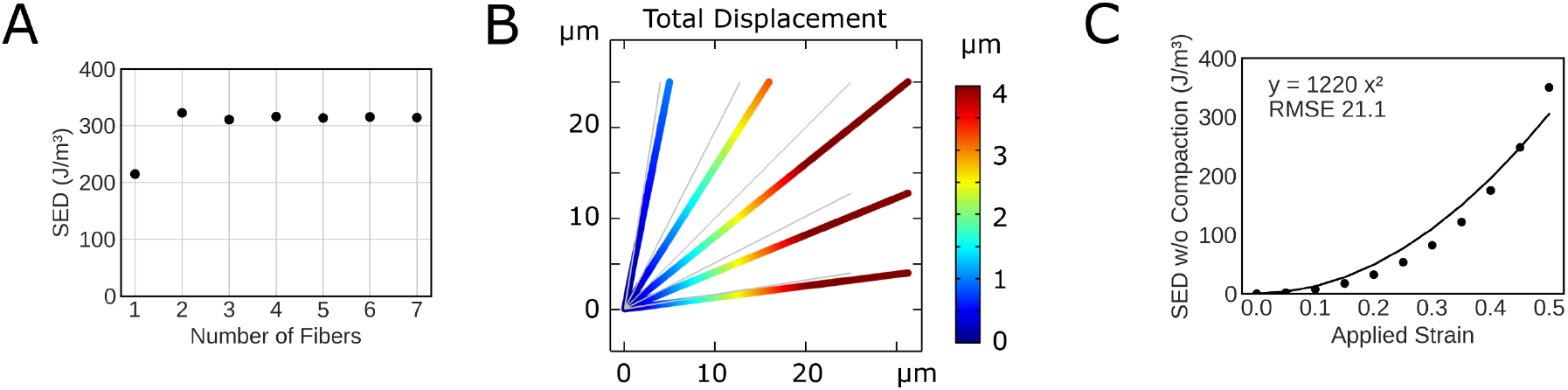
Characterization of discrete prestressed fibre model. (A) Convergence plot for strain energy density at 50% tensile strain and constant modulus (4.4 kPa), computed from a discrete fibre model consisting of different numbers of fibres; (B) Deformation of a prestressed fibre model of clot at 50% tensile strain but without any platelet contraction; (C) Strain energy density at 50% applied strain in the absence of any contractile strain shows approximately quadratic increase with externally applied strain, where RMSE = root mean square error.

**Figure S4.**
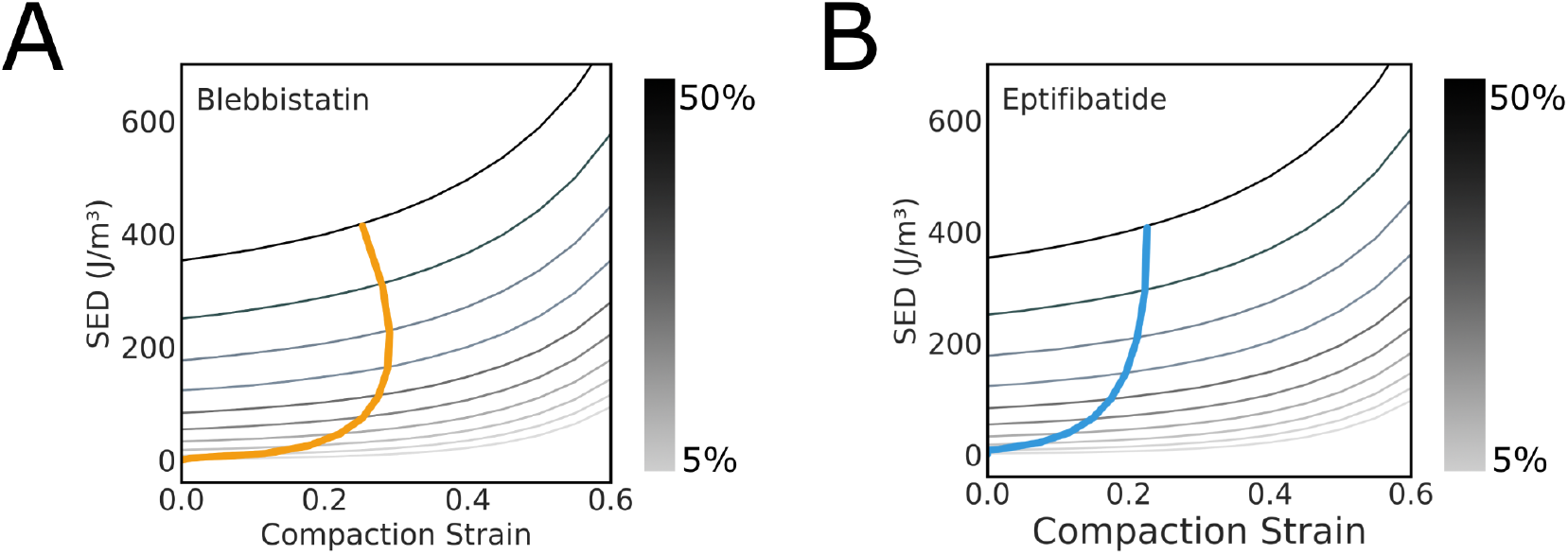
Mapping of experimental strain energy densities at applied strains from 5% to 50% onto predictions from discrete prestressed fibre model for blebbistatin (A) and eptifibatide (B).

